# Leaked genomic and mitochondrial DNA contribute to the host response to noroviruses in a STING-dependent manner

**DOI:** 10.1101/2021.08.26.457800

**Authors:** Aminu S. Jahun, Frederic Sorgeloos, Yasmin Chaudhry, Sabastine E. Arthur, Myra Hosmillo, Iliana Georgana, Rhys Izuagbe, Ian G. Goodfellow

## Abstract

The cGAS-STING pathway is central to the IFN response against DNA viruses. However, recent studies are increasingly demonstrating its role in the restriction of some RNA viruses. Here we show that the cGAS-STING pathway also contributes to the IFN response against noroviruses, positive-sense single-stranded RNA viruses that are now one of the most common causes of infectious gastroenteritis world-wide. We show a significant reduction in IFN-β induction and a corresponding increase in viral replication in norovirus-infected cells following STING inhibition, knockdown or deletion. Upstream of STING, we show that cells lacking either cGAS or IFI16 also have severely impaired IFN responses. Further, we demonstrate that immunostimulatory host genome-derived DNA, and to a lesser extent mitochondrial DNA, accumulate in the cytosol of norovirus-infected cells. And lastly, overexpression of the viral NS4 protein was sufficient to drive the accumulation of cytosolic DNA. Together, our data elucidate a role for cGAS, IFI16 and STING in the restriction of noroviruses, and demonstrate for the first time the utility of host genomic DNA as a damage-associated molecular pattern in cells infected with an RNA virus.

**Highlights:**

- cGAS, IFI16 and STING are required for a robust IFN response against noroviruses
- Nuclear and mitochondrial DNA accumulate in the cytosols of infected cells
- Viral NS4 mediates accumulation of cytosolic DNA

## Introduction

Viruses and other pathogens are detected by a myriad of receptors called pattern recognition receptors (PRRs) [1]. These receptors are present in various compartments in the host cell, including on cell surfaces (such as toll-like receptors [TLRs] and c-type lectin receptors), in endosomes (TLRs), and in the cytosol (such as the retinoic acid-inducible gene 1 [RIG-I]-like receptors, or NOD-like receptors). They are germline encoded and are able to detect pathogen-associated molecular patterns (PAMPs) which are conserved molecular patterns unique to pathogens (such as uncapped 5’ tri-phosphorylated RNA or double-stranded RNA), or common molecular signatures in aberrant conditions (such as the presence DNA in the cytosol). Activation of these sensors initiates signaling cascades that trigger the release of interferons (IFNs) and other cytokines, and promote the restriction of the invading pathogen.

Noroviruses are positive-sense single-stranded RNA viruses, with ∼7.4kb genomes comprising 3-4 open-reading frames and a poly(A) tail [2]. The human noroviruses (HuNoVs), against which there are no approved vaccines or treatment, are the commonest causes of infectious gastroenteritis worldwide, with up to 677 million cases and more than 200,000 deaths every year in children under 5 [3–5]. The murine norovirus (MNV) is broadly used as a surrogate model for studying the biology of noroviruses due to the availability of a reverse genetics system and a robust animal model, as well as an efficient virus culture system in widely available cell lines and primary cells [2]. Using MNV as a model, MDA5 and NLRP6 have been previously demonstrated to play important roles in initiating IFN responses following infection with noroviruses, both *in vivo* and in primary cells [6–8]. The distribution of these receptors is complementary and varies with cell type, such that MDA5 is largely expressed in myeloid cells, while NLRP6 is predominantly expressed in epithelial cells [8]. However, the effect of NLRP6 depletion on virus replication is only marginal [8]. Additionally, depletion of STAT1, a central mediator of signalling downstream of IFN receptors, leads to profoundly higher viral titres than those following MDA5 depletion, both *in vivo* [6,9] and *in vitro* [6,10], suggesting the presence of other receptors or pathways that contribute to innate immune responses against noroviruses.

While the cGAS-STING pathway is mostly associated with restriction of DNA viruses [1], there have been reports of its role in the restriction of replication of RNA viruses, in both IFN dependent and independent manners [11–15]. This is thought to be mediated by the presence of leaked mitochondrial DNA (mtDNA) in the cytosol resulting from viroporin-mediated calcium imbalances [12], infection-induced mitochondrial damage [11,15], or via an as yet undefined mechanism downstream of IL-1β receptor signalling [16]. This leaked mtDNA acts as a Damage-Associated Molecular Pattern (DAMP), activating host responses downstream of STING. Thus far, there are no reports of genomic DNA acting as DAMPs in the same manner, although previous studies have demonstrated STING activation following genomic DNA leakage into the cytosol in tumour cells [17,18], following extensive DNA damage from irradiation [19], or from exposure to some anticancer agents [20]. Indeed STING-dependent IFN responses to leaked genomic DNA are thought to significantly contribute to the chemotherapeutic activities of certain anticancer drugs [21,22].

In the current study, we observed a considerable attenuation of IFN responses and a commensurate increase in viral replication in norovirus-infected primary cells and cell lines following treatment with small molecule inhibitors of STING. We also show a reduction in induction of IFNs and IFN-stimulated genes (ISGs) in norovirus-infected STING^-/-^ cells, as well as cGAS^-/-^ and IFI16^-/-^ cells. We further demonstrate a substantial accumulation of nuclear DNA, and to a lesser extent mtDNA, in the cytosol of infected cells, likely mediated by the viral NS4 protein. In summary, we show for the first time a host-pathogen dynamic whereby genomic DNA acts as a damage-associated molecular pattern in cells infected with an RNA virus.

## Results

### MNV induces IFNs only in STING-competent cells

The MNV VF1 protein counteracts induction of type I IFNs through an as yet unknown mechanism [23,24]. A common and significant challenge in studying the functions of MNV proteins, including VF1, is that primary murine macrophages and MNV-permissive cell lines are often difficult to transfect [25,26], and the transfection process itself frequently leads to induction of IFNs. To circumvent these limitations, we transduced two easy-to-transfect cell lines, HeLa and HEK293T, with the MNV receptor CD300lf to make them permissive to MNV infection [27,28]. To examine whether VF1 inhibits IFN induction in human cells, the CD300lf-expressing HeLa and HEK293T cells were infected with wild-type MNV1 or the previously described VF1-negative mutant M1 in which a stop codon was introduced at position 17 of VF1 without affecting the underlying VP1 sequence [23]. As seen previously in RAW264.7 cells [23], HeLa-CD300lf cells infected with MNV1 induced substantial increase of IFN-β, with the M1 virus inducing higher levels of IFN-β compared to cells that were infected with wild type MNV1 (Figure 1a). By contrast, there was little or no IFN-β induction in HEK293T-CD300lf cells infected with either the wild type virus or M1, with no difference seen in IFN induction between cells infected with the wild type MNV1 compared to those infected with the M1 mutant (Figure 1a). These findings suggest that a factor or pathway present in HeLa and RAW264.7 cells, but absent in HEK293T is required for both a robust induction of IFN in MNV-infected cells, as well as the phenotypic differences observed between cells infected with the wild type and M1 viruses.

**Figure 1.**
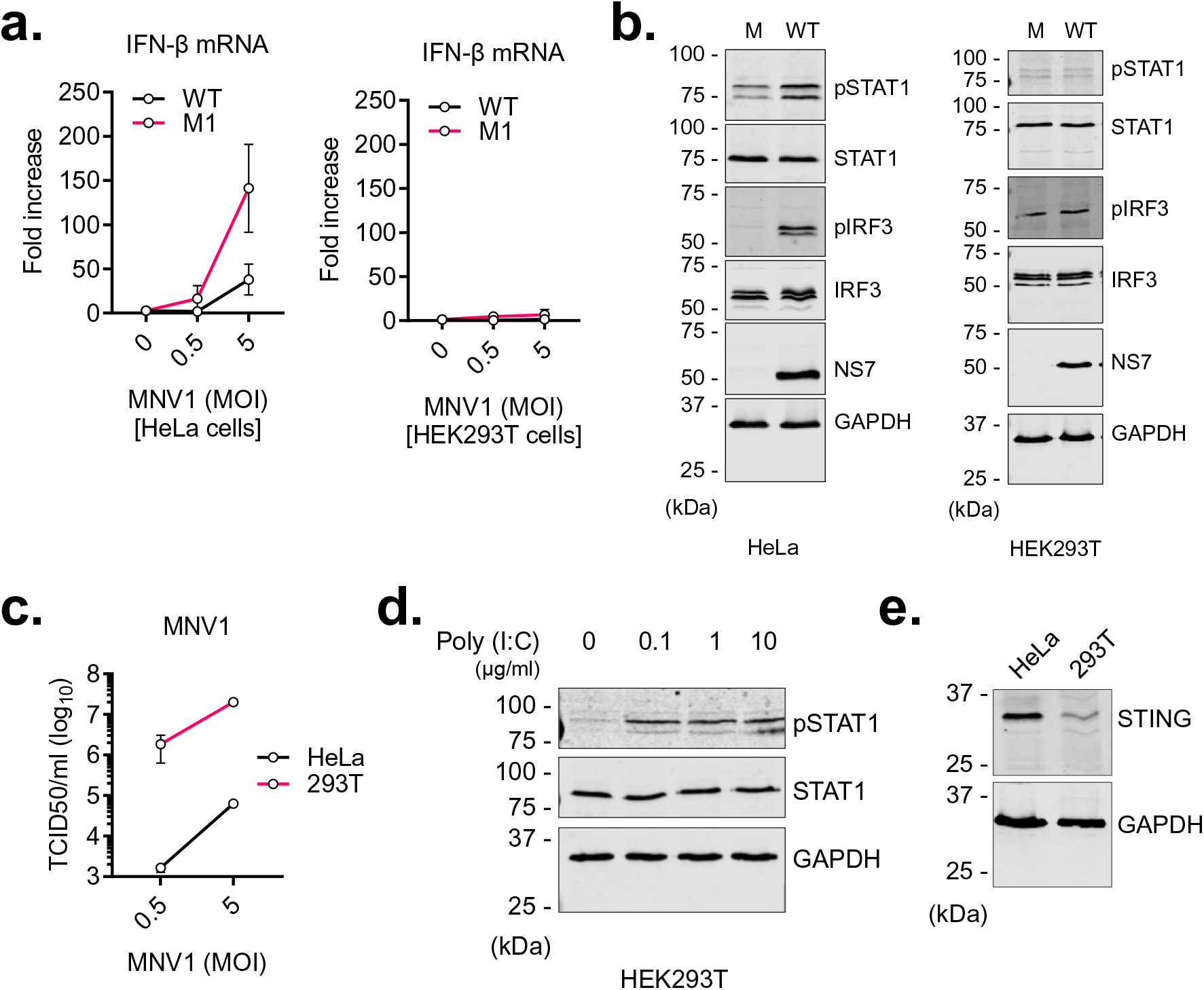
MNV induces IFNs only in STING-competent cells (a) HeLa-CD300lf and HEK293T-CD300lf cells were infected at the indicated MOIs with either wild-type MNV1 or the VF1-deleted M1 mutant. The cells were harvested 10h after infection and subjected to RT-qPCR. Data represent two independent experiments done in triplicates, and is shown relative to mock-infected cells, and normalised to human β-actin. (b) HeLa-CD300lf and HEK293T-CD300lf cells were either mock-infected, or infected at an MOI of 10 with either wild-type MNV1 or the VF1-deleted M1 mutant. The cells were harvested 10h after infection and assessed by western blotting for the indicated proteins. (c) HeLa-CD300lf and HEK293T-CD300lf cells were infected at the indicated MOIs with wild-type MNV1. The cells were harvested 10h after infection and infectious viral titres were determined using TCID50. Data represent two independent experiments, each done in triplicates. (d) HEK293T-CD300lf cells were transfected with indicated amounts of poly (I:C). The cells were harvested 6h after transfection and assessed by western blotting for the indicated proteins. (e) Lysates from HeLa-CD300lf and HEK293T-CD300lf cells were assessed by western blotting for STING and GAPDH.

To further examine IFN induction in HeLa-CD300lf and HEK293T-CD300lf cells following infection with MNV, the cells were either mock-infected, or infected with wild type MNV1 at a high MOI, harvested 10 hours post infection and the lysates assessed using western blotting. As shown in Figure 1b (right panel), there was no difference in the levels of phospho-IRF3 and phospho-STAT1 between the mock-infected cells and cells infected with MNV1 in HEK293T-CD300lf cells. This contrasts with infection in HeLa-CD300lf cells (Figure 1b, left panel) where a significant increase is seen in the levels of both phospho-IRF3 and phospho-STAT1 following infection with MNV1. We also observed profoundly higher levels in viral titres from the HEK293T-CD300lf cells compared to HeLa-CD300lf cells (Figure 1c). The HEK293T-CD300lf cells have an intact RNA sensing pathway as they are able to phosphorylate STAT1 in response to poly (I:C) transfection (Figure 1d). Both being epithelial-like cell lines, HeLa and HEK293T cells likely express the same repertoire of components of the IFN response pathway, with an exception of the adapter protein STING that is only marginally expressed in HEK293T cells, compared to HeLa cells (Burdette *et al*.[29], and Figure 1e). Taken together, these data demonstrate an attenuation of the IFN response to MNV1 infection in HEK293T-CD300lf cells, compared to HeLa-CD300lf and RAW264.7 cells, and putatively suggesting a role for STING, or other factors non-functional in HEK293T-CD300lf cells, in the IFN response against noroviruses.

### Small-molecule inhibition of STING activation enhances replication of noroviruses in cell lines and primary cells

To explore the role of STING in the antiviral response against MNV, we made use of the recently described covalent small-molecule inhibitors of STING; C-176 and H-151 [30]. As shown in Figure 2a, both molecules were able to inhibit induction of IFN-β following transfection of poly (dA:dT) in RAW264.7 cells, but not poly (I:C), consistent with previously published results. To determine whether STING is required for induction of IFNs following infection with MNV, RAW264.7 cells were pre-treated with DMSO or titrated doses of C-176 or H-151, and then infected with wild type MNV1 at a high MOI. The cells were harvested 9 hours post infection and subjected to RT-qPCR. As shown in Figure 2b, there was a significant dose-dependent decrease in IFN-β induction in cells treated with either C-176 or H-151, compared to DMSO control. These data suggest that STING is required for a robust induction of IFNs in RAW264.7 cells infected with MNV1.

**Figure 2.**
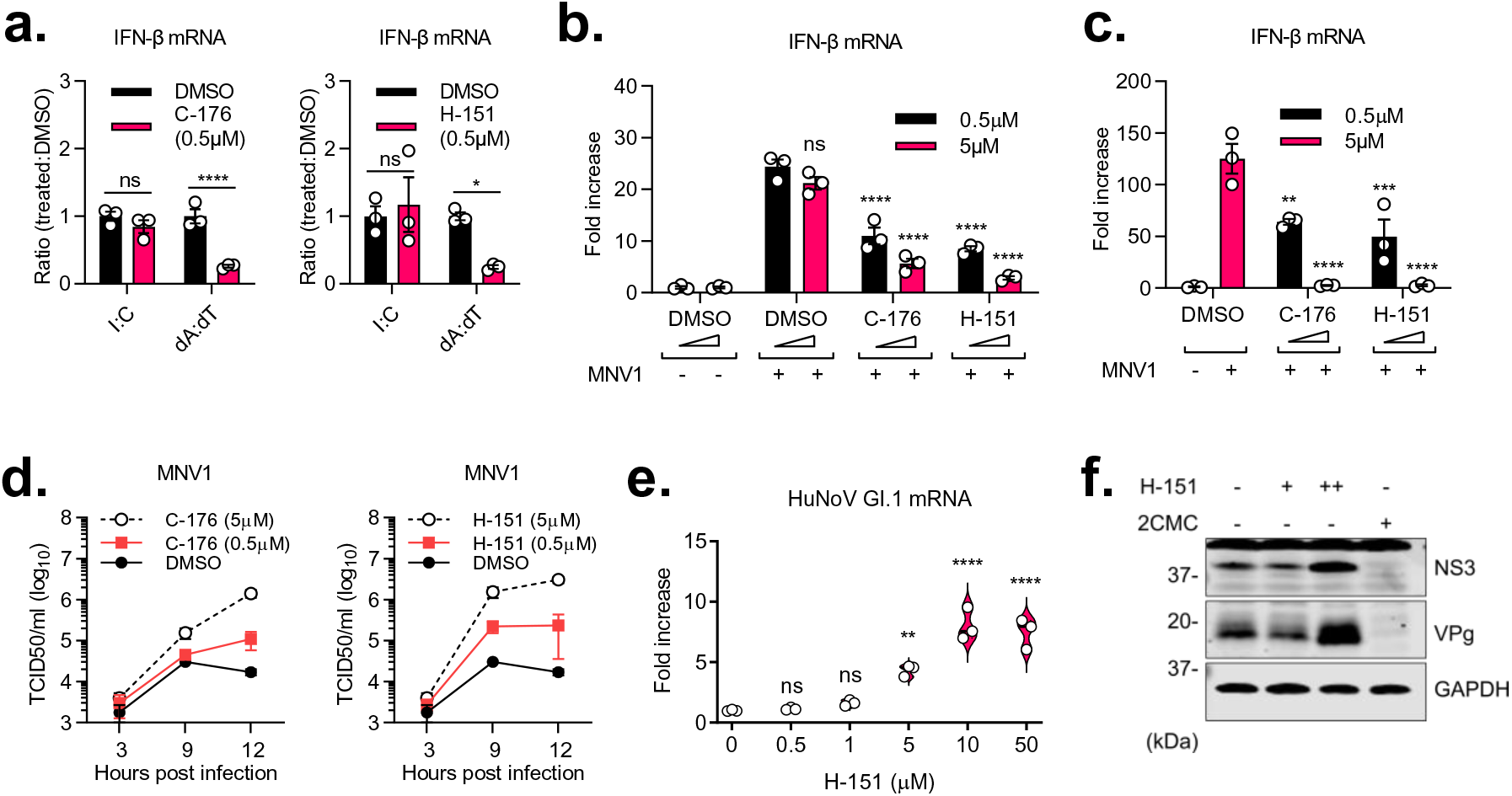
Small-molecule inhibition of STING activation enhances replication of noroviruses in cell lines and primary cells (a) RAW267.4 cells pre-treated with DMSO, 0.5μM C-176 or 0.5μM H-151 for 2h, were either mock-transfected, or transfected with 1μg poly (I:C) or poly (dA:dT). The cells were harvested after 2h and subjected to RT-qPCR. Data represent experiments done in triplicates, and is shown relative to mock-transfected cells, and normalised to mouse Gapdh (b) RAW267.4 cells pre-treated with DMSO, or indicated amounts of C-176 or H-151 for 2h, were either mock-infected or infected with wild-type MNV1 at an MOI of 10. The cells were harvested 9h post-infection and subjected to RT-qPCR. Data represent two independent experiments done in triplicates, and is shown relative to mock-infected cells, and normalised to mouse Gapdh (c) BMDM cells pre-treated with DMSO, or indicated amounts of C-176 or H-151 for 2h, were either mock-infected or infected with wild-type MNV1 at an MOI of 10. The cells were harvested 12h post-infection and subjected to RT-qPCR. Data represent two independent experiments done in triplicates, and is shown relative to mock-transfected cells, and normalised to mouse Gapdh (d) BMDM cells pre-treated with DMSO, or indicated amounts of C-176 (left panel) or H-151 (right panel) for 2h, were either mock-infected or infected with wild-type MNV1 at an MOI of 10. The samples were harvested at different time points post infection and infectious viral titres were determined using TCID50. Data represent two independent experiments, each done in triplicates. (e) Pre-seeded HGT-NV cells were treated with DMSO or indicated doses of H-151. The cells were harvested 24h post treatment and subjected to RT-qPCR. Data represent two independent experiments done in triplicates, and is shown relative to DMSO-treated cells, and normalised to human β-actin. (f) Pre-seeded HGT-NV cells were treated with DMSO, H-151 (+=0.5µM, ++=5µM), or 2CMC (200µM). The cells were harvested 72h post treatment and assessed by western blotting for the indicated proteins.

To examine the role of STING in primary macrophages, bone marrow cells from C57BL/6 mice were differentiated into bone marrow-derived macrophages (BMDMs), pre-treated with either DMSO or titrated doses of C-176 or H-151, and subsequently infected with MNV1 at a high MOI. The cells were harvested 12 hours post infection and subjected to RT-qPCR. As shown in Figure 2c, and consistent with data from assays in RAW264.7 cells, there was a significant dose-dependent decrease in IFN-β induction in MNV-infected BMDMs following treatment with the small-molecule inhibitors of STING. To determine the role of STING in restricting MNV1 replication, BMDMs were pre-treated with DMSO or titrated doses of C-176 or H-151, and then infected with MNV1. The samples were harvested at different time points post infection and infectious viral titres were determined using TCID50. As shown in figure 2d, there is a significant dose-dependent increase in viral titres following treatment with STING inhibitors. Altogether, these data indicate that STING plays an important role in the antiviral responses to MNV1 in both primary macrophages and cell lines.

Next, we wanted to see if there is a similar role for STING in the IFN responses against human noroviruses. For this, we made use of the recently described HGT-NV cells [31], a human gastric tumour cell line stably harbouring a GI.I human norovirus replicon. The cells were treated with DMSO or indicated doses of H-151, harvested 24 hours post treatment and subjected to RT-qPCR. As shown in Figure 2e, inhibition of STING activation led to a significant increase in the HuNoV genomes, in a dose-dependent manner. We also saw a marked increase in viral proteins at 72 hours after treatment in these cells (Figure 2f). Taken together, these results suggest that STING contributes to the restriction of both human and murine noroviruses during replication.

### STING^-/-^ cells induce an attenuated antiviral response against noroviruses

To confirm the role of STING in the IFN responses against noroviruses, we used the CRISPR/Cas9 system to generate two clones of STING^-/-^ RAW267.4 cell lines (Figure 3a). Both clone 14 (C14) and clone 22 (C22) STING^-/-^ cells induced IFN-β at similar levels to the wild-type cells following transfection of poly (I:C), but showed a significantly impaired response to transfected poly (dA:dT), as expected (Figure 3b). To determine the effect of STING depletion on IFN-β response to MNV, STING^+/+^ and C14 and C22 STING^-/-^ cells were either mock-infected or infected with wild-type MNV1 at an MOI of 5 and harvested at 9 hours post infection. Samples were then assessed for IFN-β mRNA by RT-qPCR. Both clones of STING^-/-^ cells showed a significant decrease in IFN-β induction and ISGs following infection with MNV1 (Figure 3c) at both high and low MOI, in agreement with data from the small-molecule inhibition experiments, and overall confirm a role for STING in the antiviral responses to MNV1 infection. These data are also consistent with data obtained from RAW264.7 cell transduced with shRNA targeting STING (Figure S1).

**Figure 3.**
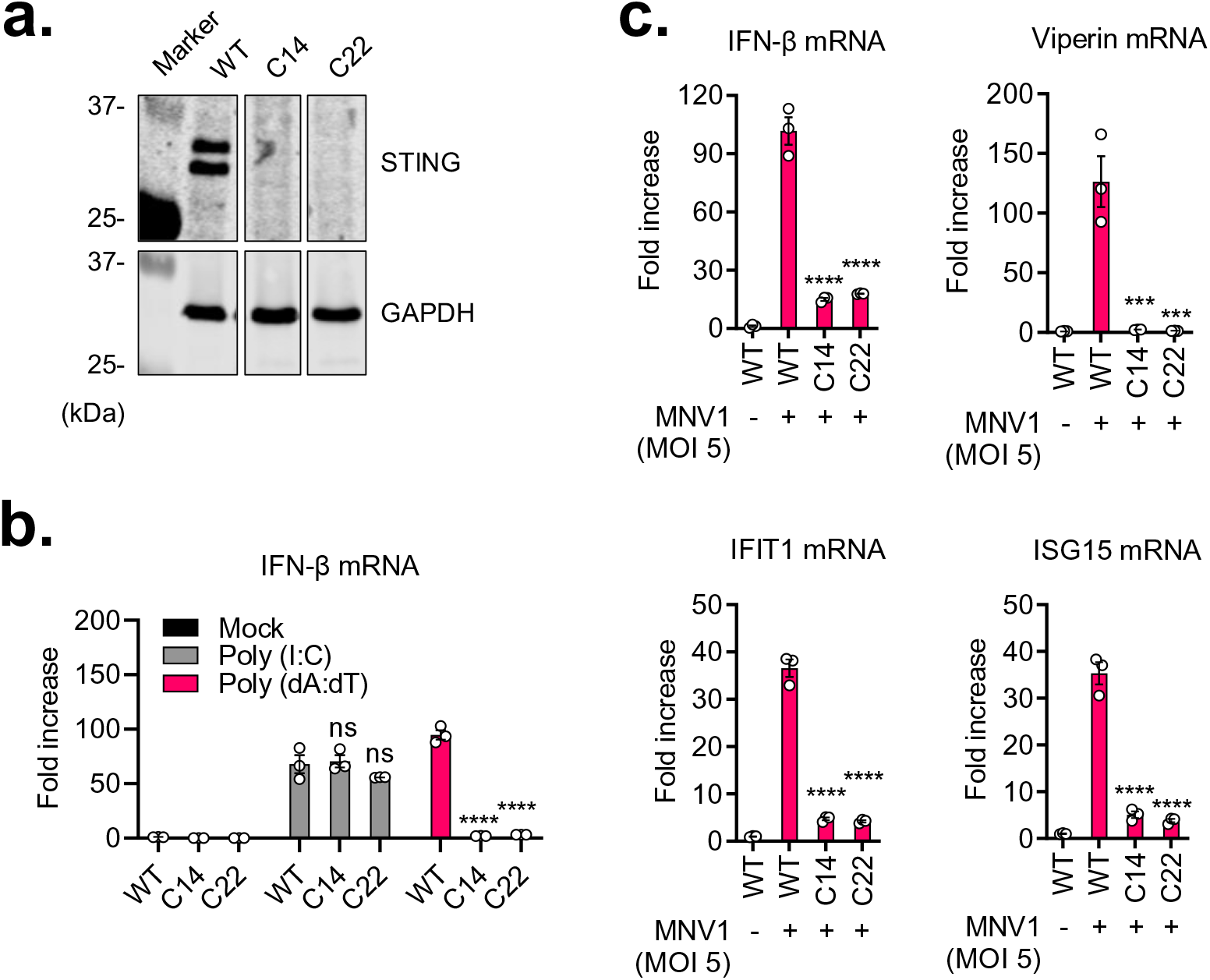
STING^-/-^ cells induce an attenuated antiviral response against noroviruses (a) STING^+/+^ (WT) and clones 14 and 22 STING^-/-^ (C14 and C22, respectively) RAW264.7 cells were assessed by western blotting for the indicated proteins. (b) STING^+/+^ (WT) and clones 14 and 22 STING^-/-^ (C14 and C22, respectively) RAW264.7 cells, were mock transfected, transfected with 1μg/ml of Poly (I:C), or with 1μg/ml of Poly (dA:dT) for 6h, and were subsequently harvested and assessed for IFN-β mRNA using RT-qPCR. Data is expressed relative to control and normalised to Gapdh. (c) STING^+/+^ (WT) and clones 14 and 22 STING^-/-^ (C14 and C22, respectively) RAW264.7 cells, were mock infected or infected with wild-type MNV1 at an MOI of 5 and harvested at 9h post infection. Samples were assessed for IFN-β mRNA via RT-qPCR. Data is presented relative to control and normalised to Gapdh.

### Both cGAS and IFI-16 contribute to IFN responses in norovirus-infected cells

Next, the upstream receptor that mediates STING activation in MNV-infected cells was examined. Activation of either cGAS or IFI16 leads to activation of STING and eventually induction of IFNs [1]. To explore the roles of these receptors in norovirus-infected cells, we used RAW264.7 cells stably expressing a secreted luciferase under the control of the ISG54 (ISRE) promoter (RAW-Lucia ISG cells). Wild-type, cGAS^-/-^ or IFI16^-/-^ (p204^-/-^) cells were infected with MNV1 at an MOI of 5. As controls, MAVS^-/-^, STING^-/-^, and MDA5^-/-^ were also infected at the same MOI. The supernatants were harvested at 18 hours post infection and analysed on a luminometer. As shown in Figures 4a and 4b, there was a significant decrease IFN-β induction in both cGAS^-/-^ and IFI16^-/-^, compared to wild-type cells, and comparable to the decrease seen in MAVS^-/-^, STING^-/-^, and MDA5^-/-^ cells. These data suggest that both cGAS and IFI16 are required for a robust induction in IFN-β in MNV-infected cells.

**Figure 4.**
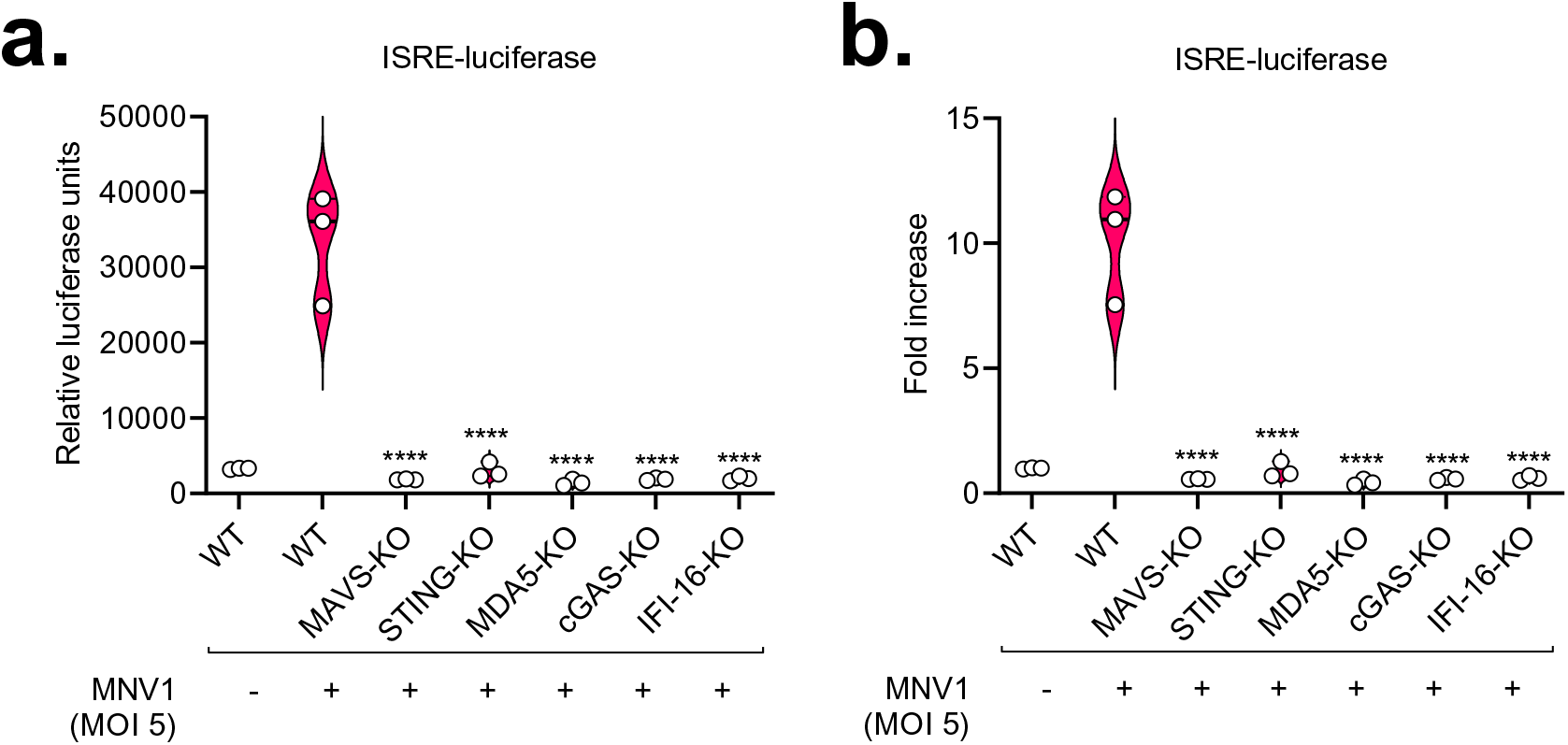
Both cGAS and IFI-16 contribute to IFN responses in norovirus-infected cells (a) and (b) Wild-type RAW-Lucia ISG cells or those with the indicated knockouts were either mock-infected or infected with wild-type MNV1 at an MOI of 5. The supernatants were harvested 18h post infection and analysed for Lucia luciferase levels on a luminometer. Data is presented as raw RLU (a) or fold increase relative to the mock-infected (b).

### Aberrant cytosolic DNA from norovirus-infected cells induces an IFN response, in a STING-dependent manner

Mitochondrial DNA has also been demonstrated to accumulate in the cytosol of host cells following infection by some RNA viruses, including Dengue virus [11], Chikungunya virus [13], Influenza A virus [12], and encephalomyocarditis virus (EMCV) [12]. This aberrant presence of DNA in the cytosol then leads to activation of the cGAS-STING pathway and the ensuing IFN response. To determine if mtDNA accumulates in the cytosol of norovirus-infected cells, RAW267.4 cells were either mock-infected or infected at an MOI of 0.5 or 5, and harvested 12 hours post-infection. The cells were then lysed in a digitonin buffer and fractionated. DNA was extracted from the cytosolic fractions and the presence of mtDNA was determined by qPCR using two previously described primer sets. As shown in Figure 5a, there was a moderate accumulation of mtDNA in the cytosol of infected cells, with higher levels observed with increased MOI. Alongside this, we also observed a sizeable dose-dependent increase in genome-derived GAPDH and HPRT DNA in the cytosol of infected cells. Taken together, these data indicate leakage of both mitochondrial and genomic DNA into the cytosol of cells infected with MNV.

**Figure 5.**
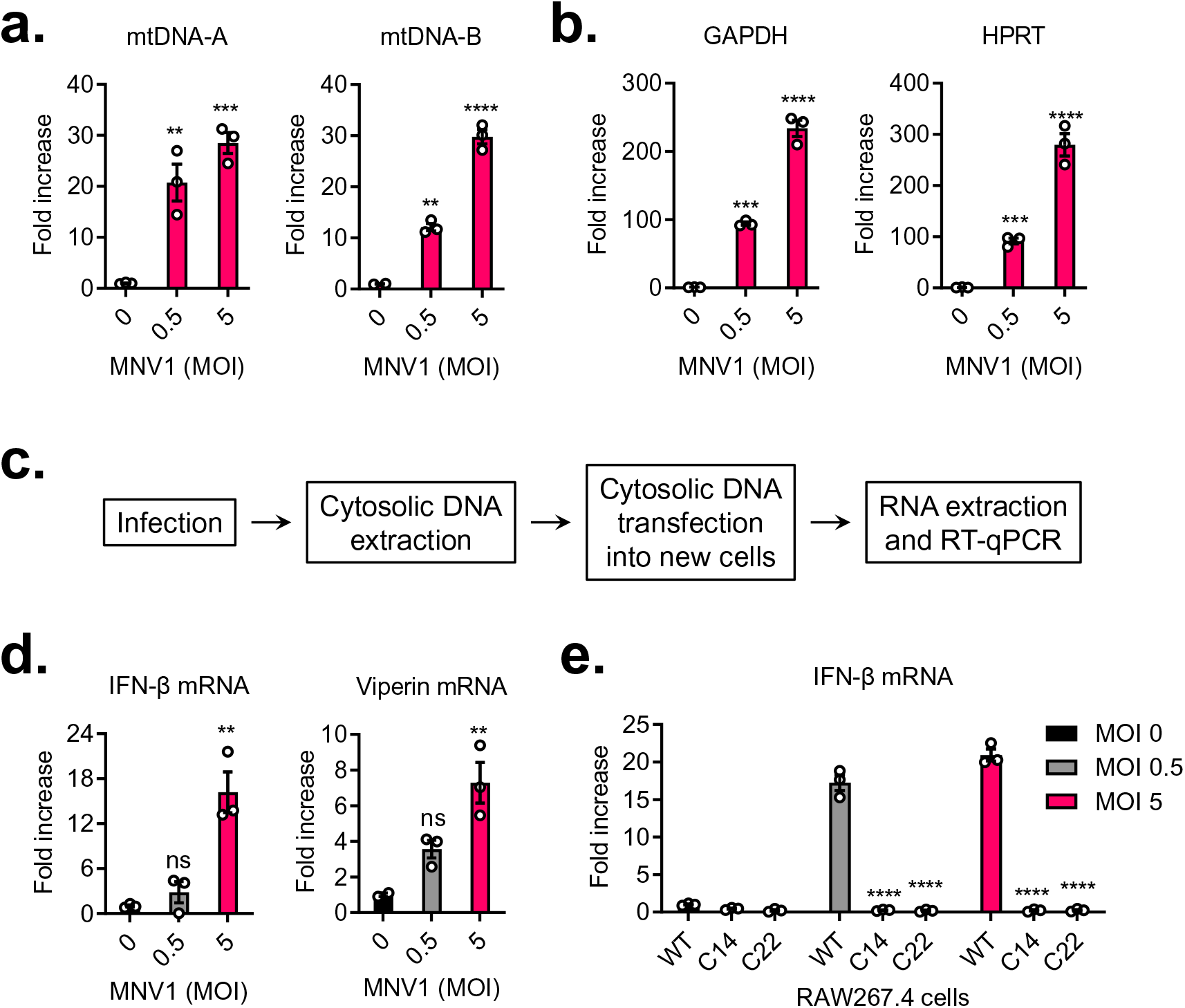
Aberrant cytosolic DNA from norovirus-infected cells induces an IFN response, in a STING- dependent manner (a) and (b) RAW264.7 cells were either mock-infected or infected with MNV1 at the indicated MOI. The cells were harvested 10h post infection and assessed for cytosolic DNA by qPCR. Data is presented relative to GFP and normalised to their corresponding whole cell fractions. (c) Schematic representation of the experiments carried out in (d) and (e) (d) and (e) RAW264.7 cells were infected with MNV1 at the indicated MOIs. The cells were harvested and fractionated 10h post infection, and DNA was extracted from the cytosolic and whole cell fractions. Pre-seeded cells were transfected with normalised amounts of the cytosolic DNA, harvested 9h post transfection, and analysed by qPCR. Data is presented relative to control and normalised to Gapdh.

To determine if this aberrant cytosolic DNA is able to induce IFNs, we transfected normalised amounts of extracted cytosolic DNA into RAW264.7 cells, and assessed for IFN-β and ISG (viperin) expression using qPCR (Figure 5c). As shown in Figure 5d, there was a significant increase in IFN-β and viperin in the cells transfected with cytosolic DNA from MNV-infected cells, compared to that from mock-infected cells, in a dose-dependent manner. This increased induction of IFN-β was seen to be STING-dependent, as there was almost no induction of IFN-β in STING^-/-^ cells transfected with cytosolic DNA (Figure 5e). Altogether, our data indicate leakage of genomic and mitochondrial DNA into the cytosol of MNV-infected cells that can activate induction of IFNs.

### The norovirus NS4 protein promotes the accumulation of cytosolic DNA

Leakage of DNA into the cytosol of cells occurs in disparate ways. For example, mtDNA leakage in A549 cells was shown to occur downstream of IL-1β signalling [16]. Treatment of these cells with IL-1β was sufficient to cause mitochondrial leakage into the cytosol, activation of IRF3 and a resultant induction of IFNs and ISGs. Since MNV infection has recently been shown to induce IL-1β secretion [32], we hypothesised that this could therefore potentially explain the leakage of mtDNA in MNV-infected cells. To determine if this is the case, RAW264.7 cells were pre-treated with an IL-1 receptor antagonist (IL-1RA), IL-1β, or both, and subsequently infected with MNV1 at an MOI of 5. The cells were harvested 12 hours post infection and analysed for IFN-β and viperin expression using qPCR. As shown in Figures S2a and S2b, we neither observed any significant decrease in IFN-β or viperin induction in the presence of IL-1RA, nor did we see any induction of IFN-β or viperin in cells treated with IL-1β. Additionally, while treatment of A549 cells with IL-1β did induce expression of IL-6, with a decrease in expression seen in cells pre-treated with IL-1RA, as expected (Figure S2c), in our hands there was no increase in IFN-β induction in cells treated with IL-1β (Figure S2d).

Next, we considered if any of the structural and non-structural viral proteins is the cause of leakage of DNA into the cytosol. One likely candidate is the NS1-2 protein that was recently shown to potentially have a viroporin activity [33]. We hypothesised that the NS1-2 protein, as a viroporin, may disrupt intracellular calcium homeostasis and eventually lead to mitochondrial leakage, similar to the M2 protein of Influenza A virus and the 2B protein of EMCV [12]. Another candidate is the NS5 protein (VPg), which was shown to be a potential interactor of PARP1 and other DNA repair enzymes in a proteomics assay [34]. Sequestration of DNA repair enzymes by VPg in the cytosol might lead to depletion of the nuclear components of these enzymes, which could then promote the leakage of genomic DNA into the cytosol. Indeed expression of the norovirus VPg in cells leads to cell cycle arrest [35,36], a phenomenon that is also seen following depletion of DNA repair enzymes. To determine if any of these is relevant to norovirus biology, we transfected plasmids encoding individual proteins of MNV1 into HEK293T-CD300lf cells. We opted to use these cells due to their high transfection efficiencies, and like RAW264.7 cells, show a significant accumulation of genomic and mitochondrial DNA in the cytosol following infection with MNV (Figures 6a and 6b). Transfected cells were harvested after 24 hours and cytosolic DNA was assessed by qPCR. A significant increase in cytosolic DNA was seen in cells expressing NS4 compared to those expressing GFP (Figure 6c). No increase in cytosolic DNA was seen in cells expressing the NS1-2 protein, VPg, or any of the other MNV proteins (Figures 6c and S3), suggesting that the NS4 protein of MNV mediates accumulation of DNA in the cytosols of MNV-infected cells.

**Figure 6.**
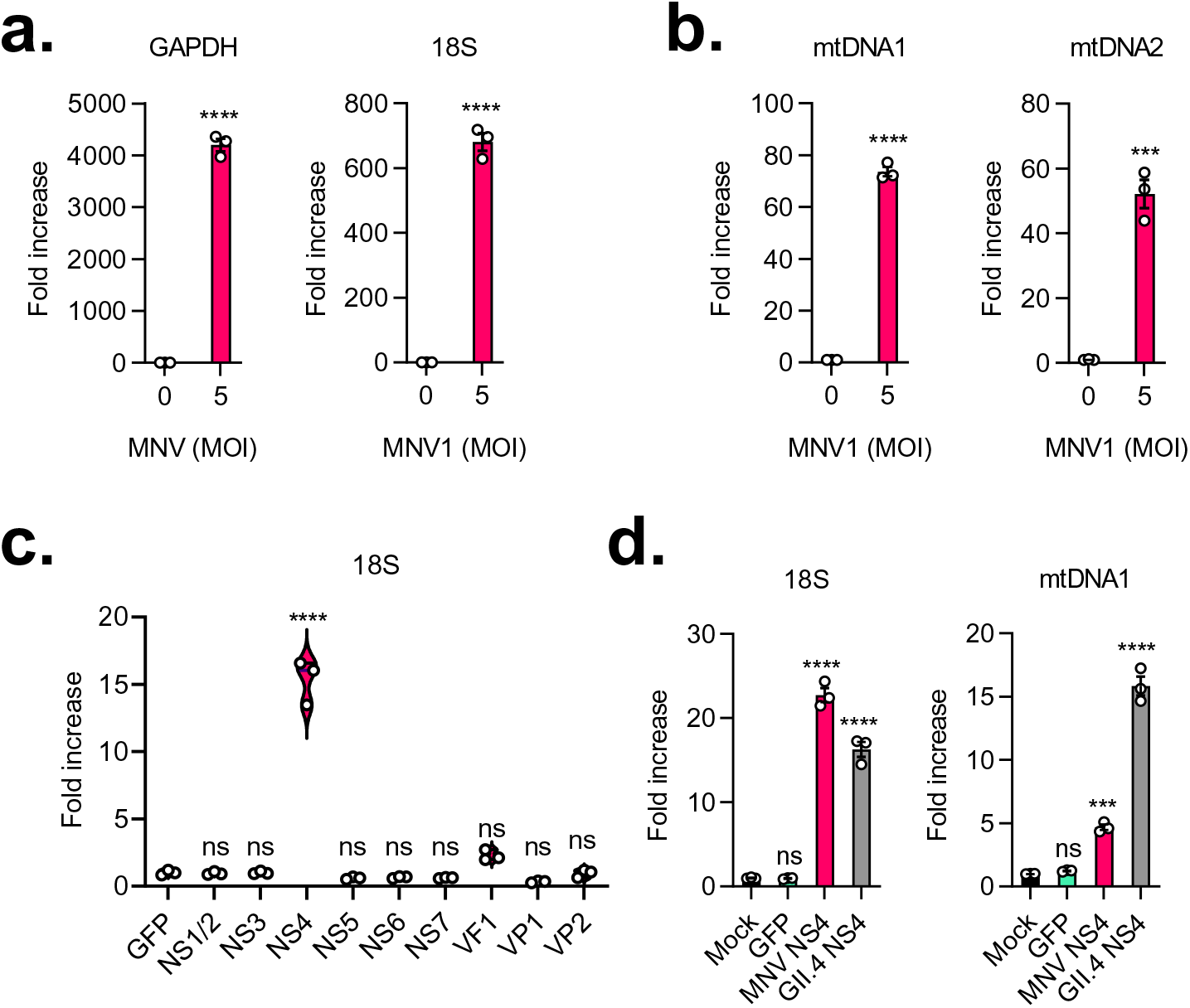
NS4 proteins of noroviruses promote accumulation of cytosolic DNA (a) and (b) HEK293T-CD300lf cells were either mock-infected or infected with MNV1 at an MOI of 5. The cells were harvested 10h post infection and assessed for cytosolic DNA by qPCR. Data is presented relative to GFP and normalised to their corresponding whole cell fractions. (d) and (d) HEK293T-CD300lf cells were transfected with Flag-tagged construct plasmids of the indicated proteins. The cells were harvested 24h post transfection and assessed for cytosolic DNA by qPCR. Data is presented relative to GFP and normalised to their corresponding whole cell fractions.

To determine if this function is conserved across the NS4 proteins of noroviruses, we explored leakage of DNA in cells expressing HuNoV NS4. For this, HEK293T-CD300lf cells were either mock-transfected, or transfected with plasmids encoding GFP, MNV1 NS4, or HuNoV GII.4 NS4. The cells were harvested 24 hours post transfection and cytosolic DNA was assessed by qPCR. While no increase in cytosolic DNA was detected in GFP-expressing cells compared to mock, as expected, a significant increase was seen in cells expressing HuNoV NS4, similar to those expressing MNV NS4 (Figure 6d). Interestingly, we observed a higher proportion of mtDNA in cells expressing HuNoV NS4 compared to those expressing MNV NS4, although levels of genomic DNA remained largely the same. Altogether, these data indicate that NS4 proteins of noroviruses are necessary and sufficient for mediating leakage of genomic and mitochondrial DNA into the cytosol.

## Discussion

Data suggesting a role for STING in restricting RNA viruses are as old as the discovery of STING itself, and the first viral proteins shown to antagonize STING function are in fact encoded by RNA viruses [15]. In this study we explored the potential role of STING in the IFN responses to norovirus infection. There was a substantial impairment of IFN responses and a corresponding increase in viral replication in norovirus-infected primary cells and cell lines following treatment with small molecule inhibitors of STING, as well as a decrease in induction of IFNs and ISGs in norovirus-infected STING^-/-^ cells, as well as cGAS^-/-^ and IFI16^-/-^ cells. Both cGAS and IFI16 can sense the presence of DNA in the cytosol [1] and activate an IFN induction signalling cascade upstream of STING. IFI16 has also been shown to be able to sense viral RNA [37], positively regulates cGAS-STING signalling [38,39], and promotes both DNA or RNA virus-induced IFN transcription in the nucleus [40]. Although STING mainly functions as an adapter protein in intracellular detection of foreign DNA, it has been shown to play important roles in the restriction of some RNA viruses through various independent mechanisms (recently reviewed by Maringer *et al.* [15], Aguirre *et al.* [41], Zevini *et al.* [42], and Ni *et al.* [43]). For example, it has been shown that STING can promote fusion-mediated IFN induction in cells infected with Influenza A virus in a manner independent of cGAS [44], and facilitate IFN induction downstream of cGAS in a manner dependent on the viral M2-mediated leakage of mtDNA [12]. In cells infected with DENV, membrane recruitment to replication complexes likely leads to leakage of mtDNA that triggers IFN induction via STING, in a process contingent on cGAS activation [11]. Recently, it was also demonstrated that STING can inhibit host and viral translation in cells infected with a wide variety of RNA viruses in a RIG-I dependent manner [14]. In addition, at least for JEV, IFN induction is largely dependent on a RIG-I/STING-dependent pathway [45]. And lastly, as were preparing this manuscript for submission, another group also published data that corroborated ours, showing a role for cGAS and STING in IFN responses against MNV, using knockout and overexpression assays [46].

The discovery of the role for the cGAS-STING pathway in the restriction of norovirus replication is significant, and potentially broadens the current MDA5/MAVS/IFN-centric understanding of the innate immune restriction of noroviruses to one that encompasses both IFN-dependent and independent pathways (Figure 7). For instance, STING can inhibit replication of RNA viruses via translation shutoff [14]. Since STING is itself an ISG [47], this could explain a previously described ability of type I IFNs to inhibit translation of MNV proteins independent of PKR [48]. STING is also involved in inflammasome activation following detection of pathogens, and this knowledge could therefore facilitate future studies of the complex relationship between noroviruses and commensal bacteria [49,50]. Importantly, this also potentially explains the discrepancy between *in vivo* and *in vitro* results from studies on IFN responses to the human norovirus. For example, studies in Huh7 and 293FT cells have shown no IFN responses to HuNoV [51,52], while human challenge studies [53], studies in animal models [54,55], and *in vitro* studies in organoids [56–58], and replicon-containing cell lines [31] have all demonstrated induction of IFNs following infection. Given that both Huh7 and 293FT cells have impaired cGAS-STING pathways [29,59,60], our data demonstrating a role of this pathway in restriction of noroviruses harmonises these various otherwise conflicting data.

**Figure 7.**
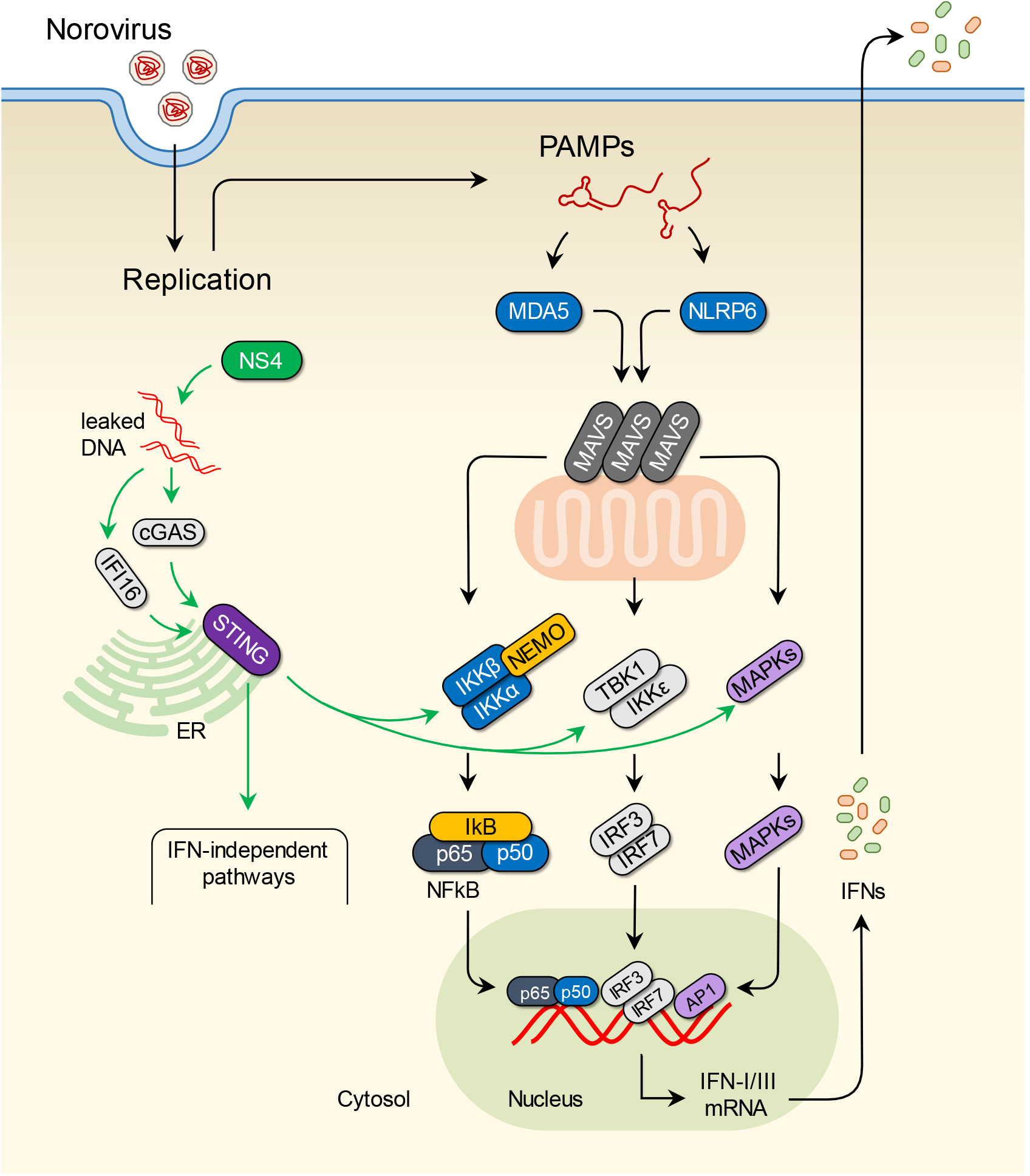
Contribution of leaked genomic and mitochondrial DNA to the host response to noroviruses in a STING-dependent manner. The MDA5/NLRP6/MAVS pathway is thought to play a central role in the detection of PAMPs in norovirus-infected cells, initiating a signalling cascade that leads to induction of type I and type III IFNs. Our data show that accumulation of genomic and mitochondrial DNA in the cytosol, driven by the viral NS4, likely activates the cGAS/IFI16/STING pathway. The combined activation of the two pathways is required for a robust IFN response and restriction of norovirus replication.

We considered if the NS1-2 protein was the cause of leakage of DNA into the cytosol given that it has been suggested to be a viroporin [33]. The M2 protein of Influenza A virus and the 2B protein of EMCV have been shown to mediate leakage of mtDNA by upsetting the intracellular calcium balance in a manner dependent on their viroporin activity [12]. We also considered the NS5 protein (VPg), a potential interactor of PARP1 and other DNA repair enzymes [34]. We hypothesized that sequestration of DNA repair enzymes by VPg in the cytosol may perhaps lead to depletion of the nuclear components of these enzymes, which in turn could lead to leakage of genomic DNA into the cytosol. This hypothesis was supported by previous studies showing that expression of the norovirus VPg in cells leads to cell cycle arrest via a yet unknown mechanism [35,36,61], given that cell cycle arrest can be a consequence of compromised DNA repair. However, our data has shown that none of these proteins induce accumulation of cytosolic DNA independently. A third hypothesis we considered for the mechanism of DNA leakage into the cytosol of infected cells was IL-1β signalling. Leakage of immunostimulatory mtDNA in West Nile virus-infected cells was previously shown to likely occur downstream of IL-1β signalling via an unknown mechanism [16,62]. Since MNV was recently shown to induce significant release of IL-1β *in vivo* [32], we considered if this pathway was also activated in norovirus-infected cells. However, we were not able to demonstrate mtDNA leakage in cells treated with IL-1β, and treatment with an IL-1RA did not affect IFN induction in MNV-infected cells.

Our finding that infection promotes leakage of genomic DNA is surprising, but not unexpected, given the substantial widespread membrane reorganisation that occurs in infected cells during the formation of virus replication complexes [63,64]. We have shown that expression of the viral NS4 protein is sufficient to cause this accumulation of DNA in the cytosol. Interestingly, a previous study demonstrated that the NS4 protein is uniquely able to form membranous complexes when overexpressed in cells [64]. Whether the ability of NS4 to promote accumulation of cytosolic DNA is related to its membrane recruitment function remains to be tested. Indeed the presence of nuclear envelope markers on norovirus replication complexes has not been previously explored. While leakage of genomic DNA into the cytosol of cells infected with RNA viruses has not been demonstrated until now, it is seen in cancer cells [17,18] and following irradiation [19] or exposure to chemotherapeutic agents such as etoposide [20]. Indeed, the efficacy of some anticancer drugs has been shown to be dependent on their ability to activate STING in this manner [21,22,65]. Given the widespread reorganisation of host cell architecture during viral replication, we expect that many more RNA viruses likely trigger leakage of genomic DNA into the cytosol.

Several RNA viruses encode proteins that counteract STING-dependent host antiviral responses [15]. The Dengue virus NS2B and Chikungunya virus capsid proteins promote degradation of cGAS in infected cells, for example [11,13]. The Influenza A virus NS1 protein binds to mtDNA and renders it less immunostimulatory [12]. Further work is required to determine if noroviruses have mechanisms to counteract or avoid this pathway. One potential target could be the GTPase-activating protein SH3 domain-binding protein 1 (G3BP1). Our group and others have recently shown that G3BP1 is sequestered within replication complexes in infected cells [34,66,67]. Since it was also recently shown to contribute to DNA sensing by cGAS [68–70], it is entirely possible that its sequestration in replication complexes also dampens its contribution to cGAS activation. Further work is however required to determine if this is the case.

In this work, we have shown a role for cGAS, IFI16 and STING in the restriction of noroviruses, and demonstrated for the first time the role of the host genomic DNA as a damage-associated molecular pattern in cells infected with an RNA virus. We have demonstrated accumulation of nuclear DNA, and to a lesser extent mtDNA, in the cytosol of infected cells, likely driven by the viral NS4 protein. Further work is required to determine the exact mechanism of this DNA leakage, as well as potential mechanisms of evasion by the viruses.

## Materials and methods

### Cells

RAW264.7, BV2, HEK293T, and HeLa cells were maintained at 37°C in complete Dulbecco’s Modified Eagle Medium (DMEM, Sigma Aldrich) containing 4500mg/ml glucose, sodium bicarbonate, and sodium pyruvate, and supplemented with 10% heat-inactivated Fetal Bovine Serum (HyClone), 10U/ml of penicillin, 100 μg/ml of streptomycin, 2mM L-glutamine (Sigma Aldrich), and non-essential amino acids (Sigma Aldrich). HEK293T and HeLa cells used in this work were generously provided by Dr Susanna M. Colaco (University of Cambridge). BSR-T7 cells, a kind gift from Karl-Klaus Conzelmann (Ludwid Maximillians University, Munich) derived from Baby Hamster Kidney (BHK) cells and expressing the T7 RNA polymerase, were maintained in complete DMEM supplemented with 0.5 mg/ml G418 (Invivogen). RAW-Lucia ISG wild-type (Invivogen, rawl-isg), MAVS-KO (Invivogen, rawl-komavs), STING-KO (Invivogen, rawl-kostg), MDA5-KO (Invivogen, rawl-komda5), cGAS-KO (Invivogen, rawl-kocgas), and IFI16-KO (Invivogen, rawl-koif16) were purchased from Invivogen.

Bone marrow-derived macrophages (BMDMs) were differentiated from bone marrow cells of C57BL/6 mice as previously described [71]. Briefly, bone marrow cells were seeded on non-treated culture plates in complete DMEM supplemented with 10% CMG14 culture supernatant which contains M-CSF. Fresh medium was added every 3 days and cells were harvested and used for experiments on day 9 or 10. This work was carried out in accordance with regulations of The Animals (Scientific Procedures) Act 1986 [72] and the ARRIVE guidelines [73]. All procedures were approved by the University of Cambridge Animal Welfare and Ethical Review Body (AWERB) and the UK Home Office and carried out under the Home Office project licence PPL 70/7689.

### Plasmids

Plasmids used in this work are listed in Table 1.

**Table 1:**
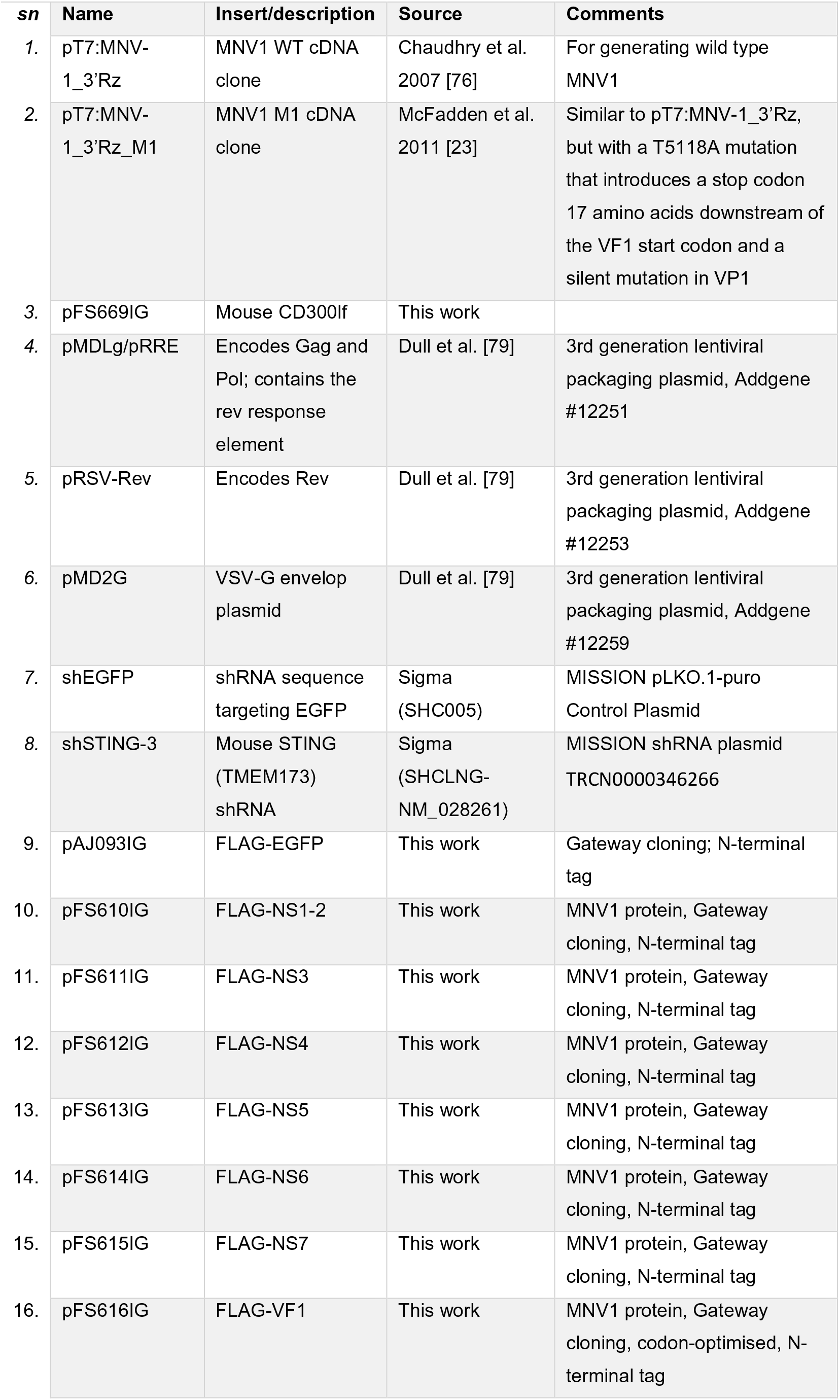
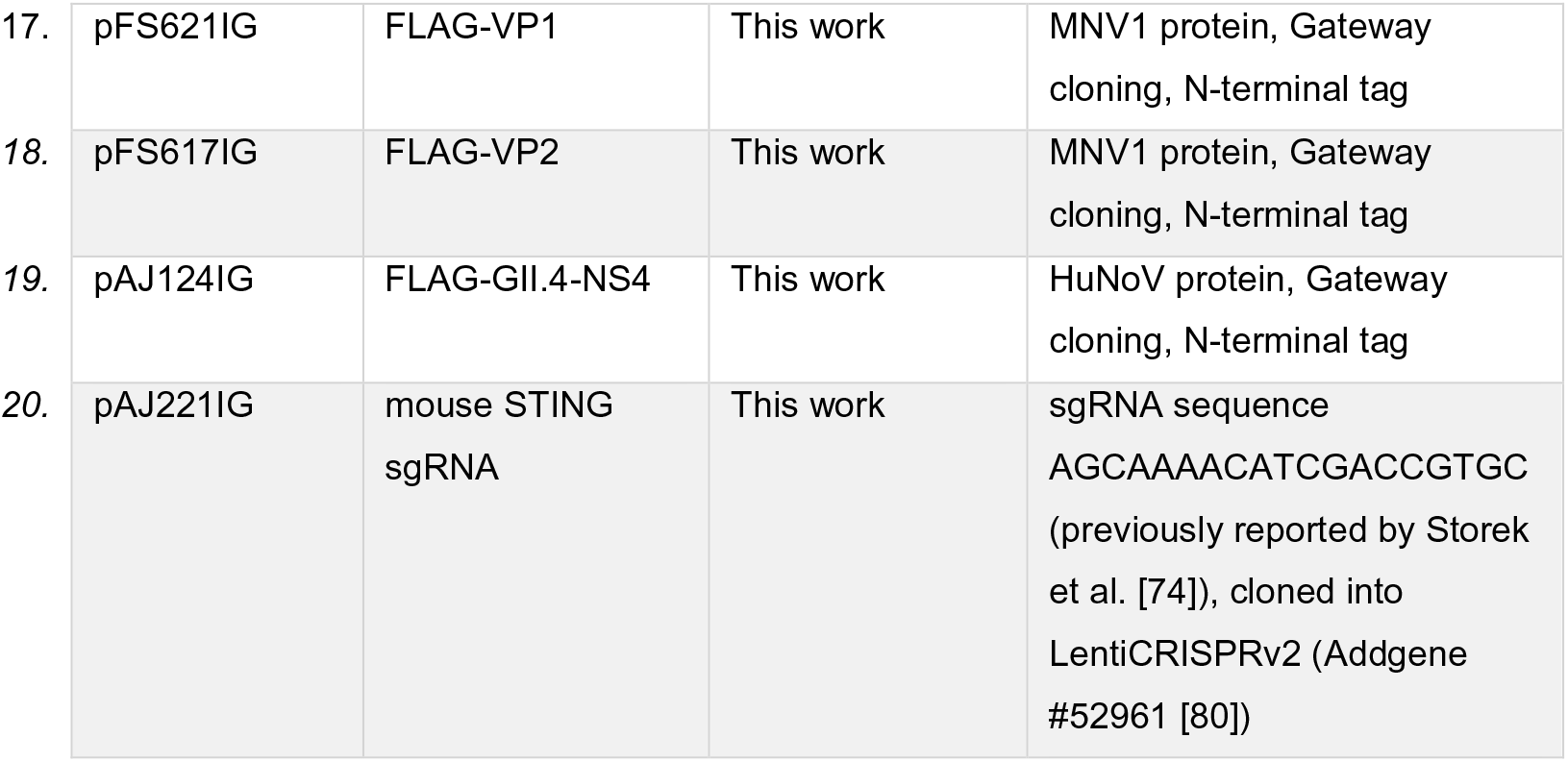
List of plasmids used.

### Lentivirus transduction

For shRNA transduction, Mission shRNA plasmids (Sigma Aldrich) were transfected together with pMDLg/pRRE, pRSV-Rev, and pMD2.G plasmids into HEK293T cells using Lipofectamine 2000. Pooled lentiviral supernatants harvested on days 2 and 3 were used to infect RAW264.7 cells. Puromycin (Invitrogen) selection was started 72 hours post-infection. The cells were cultured in 2 μg/ml puromycin until all the control cells were dead and were then maintained in 5 μg/ml puromycin.

For CD300lf lentiviral transduction, the pFS669IG plasmid encoding the mouse CD300lf was transfected together with pMDLg/pRRE, pRSV-Rev, and pMD2.G plasmids into HEK293T cells using Lipofectamine 2000. Pooled lentiviral supernatants harvested on days 2 and 3 were used to infect HeLa and HEK293T cells. CD300lf-transduced HeLa and HEK293T cells were subsequently selected using 100 μg/ml Hygromycin (Invitrogen), starting 72 hours post-infection.

For generation of STING^-/-^ RAW264.7 cells, the LentiCRISPRv2-based plasmid pAJ221IG, co-encoding the Cas9 nuclease and single guide RNA targeting mouse STING (sequence: AGCAAAACATCGACCGTGC, from Storek et al. [74]), was transfected together with pMDLg/pRRE, pRSV-Rev, and pMD2.G plasmids into HEK293T cells using Lipofectamine 2000. Pooled lentiviral supernatants harvested on days 2 and 3 were used to infect RAW264.7 cells. Puromycin (Invitrogen) selection was started 72 hours post-infection. The cells were cultured in 2 μg/ml puromycin until all the control cells were dead and were then maintained in 5 μg/ml puromycin. Single cells were FACS sorted into individual wells of 96-well plates containing conditioned media by the NIHR Cambridge BRC Cell Phenotyping Hub. Clones of cells were screened 6 weeks later by western blotting.

### Reverse genetics

The MNV1 virus was prepared via reverse genetics as previously described [75,76]. Briefly, 1.5×10^6^ BSR-T7 cells were seeded in a 6-well plate and incubated at 37°C for 3 hours. The cells were then infected with Fowlpox virus (FPV)-T7 at an MOI of 0.5 pfu/cell and incubated at 37°C for 2 hours. Then, 1μg of the pT7:MNV-1_3’Rz or the pT7:MNV-1_3’Rz_M1 MNV cDNA clones (for the wild type or VF1-deficient M1 mutant, respectively) were transfected using Lipofectamine 2000, according to the manufacturer’s instructions. The plate was incubated at 37°C for 2 days, freeze-thawed once (at −80°C overnight or longer), and titred by TCID50.

### TCID50

TCID50 by cytopathic effect (CPE) was carried out as previously described [75]. Briefly, 1:10 serial dilutions of the virus preparations were made in cell culture media and aliquoted into wells of a 96-well plate, each in 4 replicates of 50μl. Then, 2×10^4^ BV2 cells in 100μl of cell culture media was added to each well and the plate was incubated at 37°C for 5 days. The cells were subsequently assessed for CPE, and TCID50/ml was calculated using the Spearman & Kärber algorithm [77].

### Cell stimulation

Poly (I:C) (P1530, Sigma) and poly (dA:dT) (P1537, Sigma) transfections were carried out on cells pre-seeded overnight in 24-well plates using Lipofectamine 2000 (Invitrogen), according to the manufacturer’s protocol. Animal-free recombinant IL-1RA (Peprotech, AF-200-01RA), mouse IL-1β (Peprotech, AF-211-11B) and human IL-1β (Peprotech, AF-200-01B) were used for the experiments in Figure S2.

### MNV infection

Cells were incubated with the virus inoculum at the appropriate MOI on an end-to-end rotor at 37°C for an hour, then washed twice with fresh media, transferred to appropriate culture plates, and incubated at 37°C.

### Small molecule inhibition of STING

For experiments involving STING inhibition, cells were pre-treated in DMSO (Sigma), C-176 (Focus Biomolecules), or H-151 (Focus Biomolecules) for 2 hours before infection or transfection with poly (I:C) or poly (dA:dT), and the drugs are supplemented in the media onwards until the cells were harvested for end-point assays.

### Luciferase assay

RAW-Lucia ISG cells were infected as described. Clarified supernatants were harvested at 18 hours post infection, mixed with the Quanti-luc Gold reagent (Invivogen, rep-qlcg1) in a 1:1 ratio, and analysed on a Glomax Navigator Microplate Luminometer (Promega).

### Western blotting

Cells were washed in ice-cold PBS twice, resuspended in RIPA buffer (150mM NaCl, 1% NP-40, 0.5% Na deoxycholate, 0.1% SDS, 25mM Tris-HCl pH 7.4) supplemented with a protease inhibitor cocktail (and a phosphatase inhibitor cocktail when phospho-proteins were of interest), and kept on ice for 20 minutes. The sample was pipetted up and down several times and was centrifuged immediately at 10,000 x g for 10 minutes at 4°C. The supernatant was transferred to a new tube and the pellet was discarded. The sample was quantified using the BCA assay (Thermo Scientific) according to the manufacturer’s recommendations. The sample was then mixed with SDS polyacrylamide gel electrophoresis (PAGE) sample buffer (2% SDS, 10% glycerol, 0.002% bromophenol blue, 0.0625M Tris-Cl pH 6.8, 5% 2-mercaptoethanol), heated at 95°C for 5 minutes, and kept at −20°C or used immediately for SDS PAGE. Transfers were made onto 0.45μm nitrocellulose membranes. The membranes were blocked in 5% milk PBST for 1 hour at room temperature, and the primary and secondary antibodies were incubated at 4°C overnight and 1hour at room temperature respectively, with three 5-minute washes in between incubations. The membranes were subsequently scanned on an Odyssey CLx imager (LI-COR) and the results were analysed using the Image Studio Lite software version 5.2.5 (LI-COR). Antibodies used in this work are listed in Table 2.

**Table 2:** List of antibodies used for western blots.

| **sn** | **Target** | **Supplier** | **Catalogue Number** | **Dilution** |
| --- | --- | --- | --- | --- |
| 1. | Mouse GAPDH | Ambion | AM4300 | 1:20000 |
| 2. | Human GAPDH | Protein Tech | 10494-1-AP | 1:20000 |
| 3. | Human IRF3 | Abclonal | A11373 | 1:250 |
| 4. | Human pIRF3 | Abcam | ab138449 | 1:500 |
| 5. | STAT1 | Abcam | ab92506 | 1:500 |
| 6. | pSTAT1 | Cell signalling | 9167S | 1:500 |
| 7. | STING (D2P2F) | Cell signalling | #13647 | 1:500 |
| 8. | NS3 | Non-commercial | - | 1:500 |
| 9. | VPg | Non-commercial | - | 1:500 |
| 10. | NS7 | Non-commercial | - | 1:500 |

### Relative RT-qPCR

RNA extraction with on-column DNAse treatment were done using the GenElute Mammalian Total RNA Miniprep Kit (Sigma-Aldrich) according to the manufacturer’s protocol. cDNA was synthesized using the M-MLV Reverse Transcriptase (Promega), according to the manufacturer’s protocol. qPCR was carried out using a 2X SYBR Green mastermix containing 2.5mM MgCl_2_, 400μM dNTPs, 1/10,000 SybrGreen (Molecular Probes), 1M Betaine (Sigma), 0.05U/μl of Gold Star polymerase (Eurogentec), 1/5 10X Reaction buffer (750 mM Tris-HCl pH 8.8, 200 mM [NH4]_2_SO_4_, 0.1 % [v/v] Tween 20, Without MgCl_2_), and ROX Passive Reference buffer (Eurogentec), and ran on a ViiA 7 Real-Time PCR System (ThermoFisher Scientific), with a 15-second 95°C denaturation step and a 1-minute 60°C annealing/extension step for 40 cycles. Relative gene expression was calculated using the Livak method (ΔΔCt) relative to mock-transfected conditions [78], and normalized to a house keeping gene (*Gapdh* for all the mouse samples, and β-actin for the human samples). Primers used in this work are listed in Table 3.

**Table 3:**
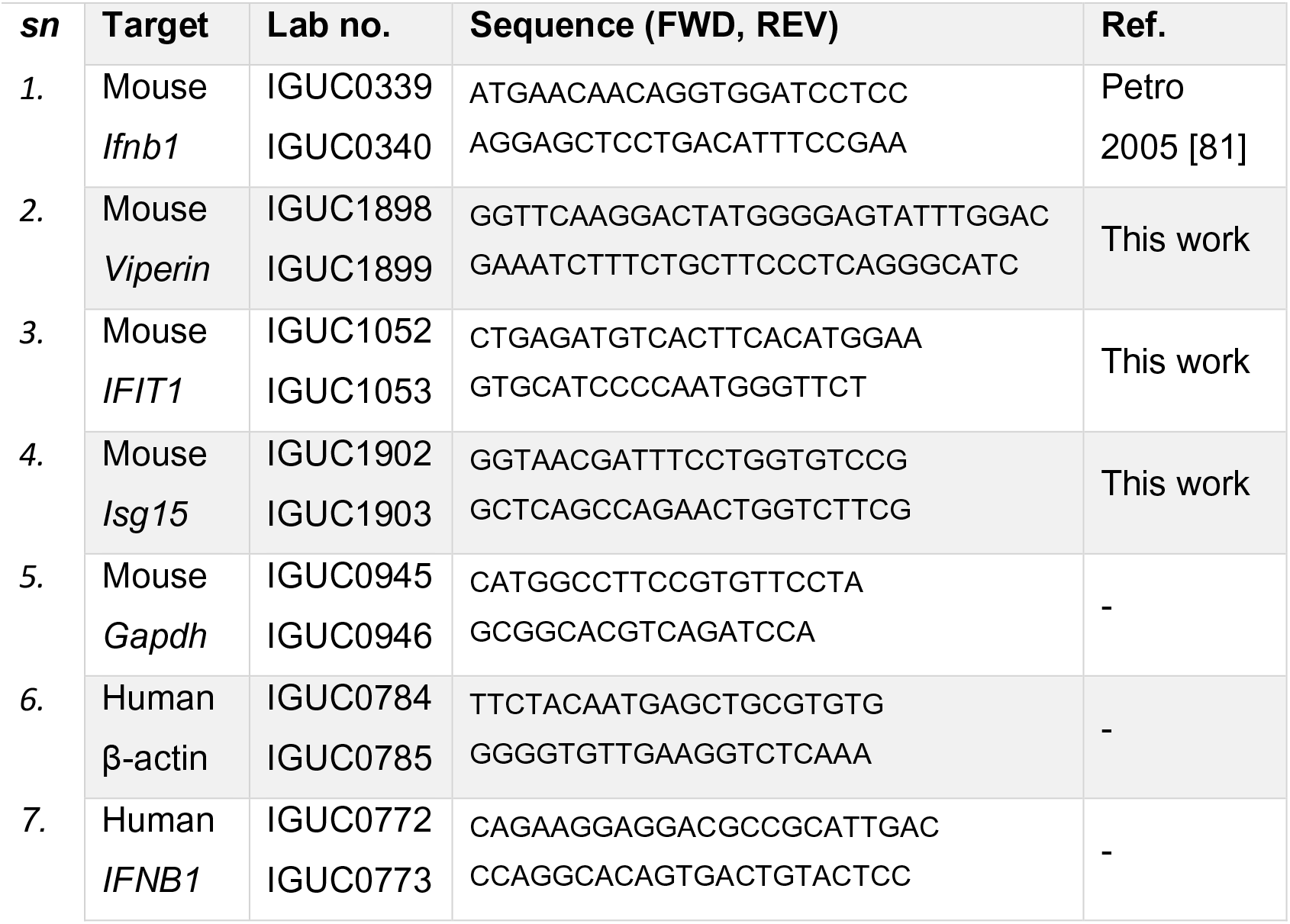
List of RT-qPCR Primers used.

### Cell fractionation and cytosolic DNA assessment

Cell fractionation for cytosolic DNA assessment was carried out using a protocol modified from Moriyama et al. [12]. Briefly, the cells were washed in 1 ml of ice-cold PBS. Cells were then resuspended in 600μl digitonin lysis buffer (150 mM NaCl, 50 mM HEPES pH 7.4, and 20 μg/ml digitonin), out of which 100μl was set aside for analysis by western blot, and 100μl was set aside as the whole cell fraction. The remaining 400μl was centrifuged at 1,000 x g for 3 minutes, and the supernatant transferred to a new tube. This was repeated twice, and then the supernatant was centrifuged at 17,000 x g for 10 minutes and transferred into a new tube. An RNAse digestion was carried out on this cytosolic fraction and the whole cell fraction by adding 2.5 μl each (1.25 units) of the RNase Cocktail Enzyme Mix (Invitrogen, AM2286) and incubated at 37°C for 2 hours. DNA was then extracted from the cytosolic fraction using the QIAquick Nucleotide Removal kit (QIAGEN) and from the whole cell fraction using the QIAamp DNA Mini Kit (QIAGEN). Both cytosolic and whole cell fractions were eluted in 100μl and diluted 1:10 in water before assessment by qPCR. Primers used for qPCR are listed in Table 4.

**Table 4:**
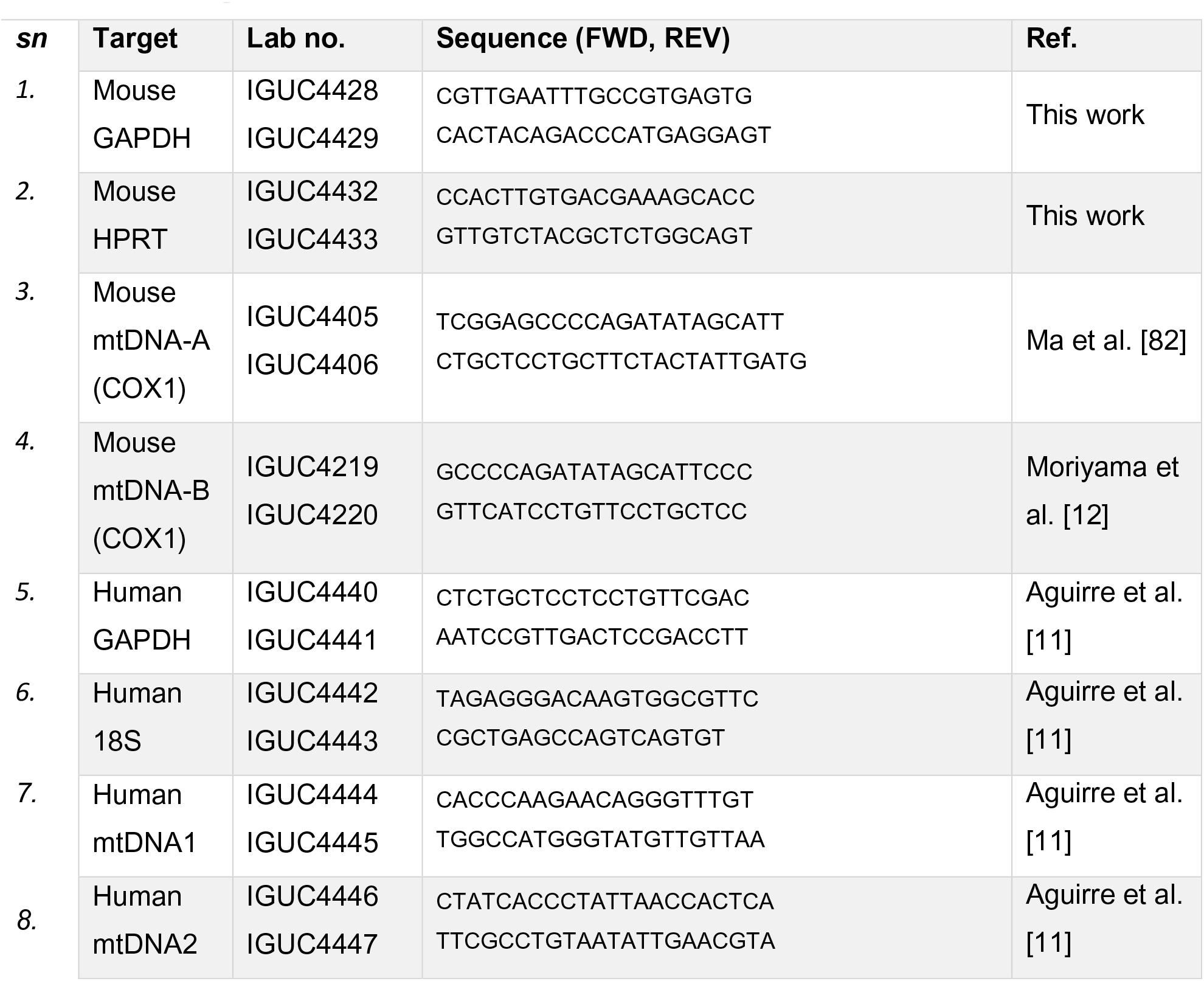
List of genomic and mitochondrial DNA Primers used.

### Statistical analysis and software

Prism 6.0 (Graph Pad) was used for all statistical analysis, and one-way repeated measures ANOVA with Bonferroni multiple comparisons tests was applied to determine statistical significance, unless where indicated otherwise. In all cases, ‘ns’, *, **, ***, and **** are used to denote p>0.05, p≤0.05, p≤0.01, p≤0.001 and p≤0.0001 respectively. The Image J software was used for all confocal micrograph preparation and Image Studio Lite 5.2 was used for western blot quantification. Clustal Omega was used for all sequence alignments, and Snapgene 4.2 was used for primer design and cloning strategies.

## Author contributions

ASJ, FS and IGG were involved in the conceptualization of the project. IGG secured funding for and supervised the work. ASJ, FS, YC, SEA, MH, IG, and RI designed and conducted the experiments, and analysed the results. ASJ wrote the initial draft of the manuscript. All authors were involved in the interpretation of the results and writing of the manuscript.

## Acknowledgements

We thank the NIHR Cambridge BRC Cell Phenotyping Hub for providing cell sorting services.

---

**Figure S1.**
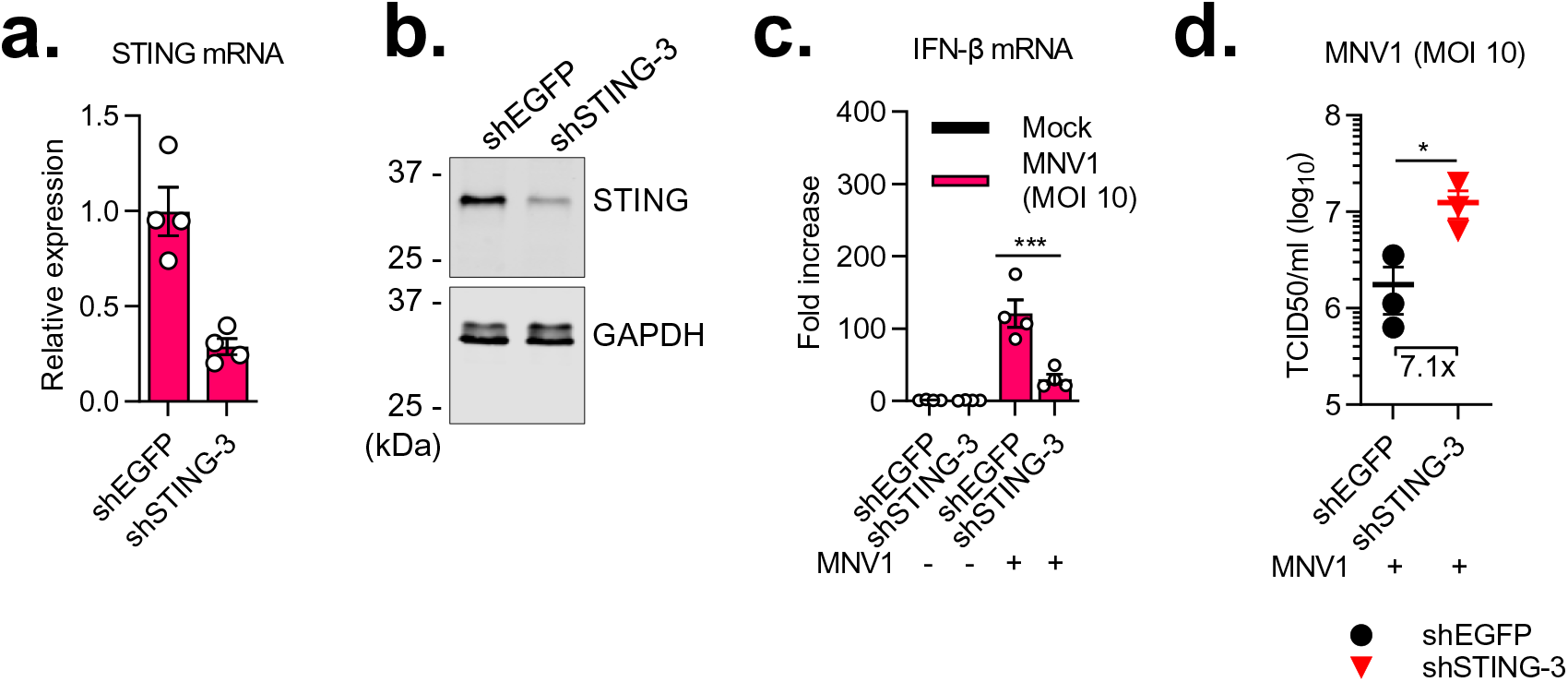
STING knockdown impairs IFN induction and increases virus replication in MNV-infected cells (a) and (b) RAW264.7 cells stably transduced with control shRNA (shEGFP) or shRNA targeting mouse STING (shSTING-3) were lysed and assessed for STING mRNA via RT-qPCR (a) or analysed via western blotting (b). Data in the left panel is presented relative to control and normalised to Gapdh. (c) and (d) RAW264.7 cells stably transduced with control shRNA (shEGFP) or shRNA targeting mouse STING (shSTING-3), were mock infected or infected with wild-type MNV1 at an MOI of 10 and harvested at 9h post infection. Samples were assessed for IFN-β mRNA via RT-qPCR (c), or infectious viral titres were determined using TCID50 (d). Data in (c) is presented relative to control and normalised to Gapdh.

**Figure S2.**
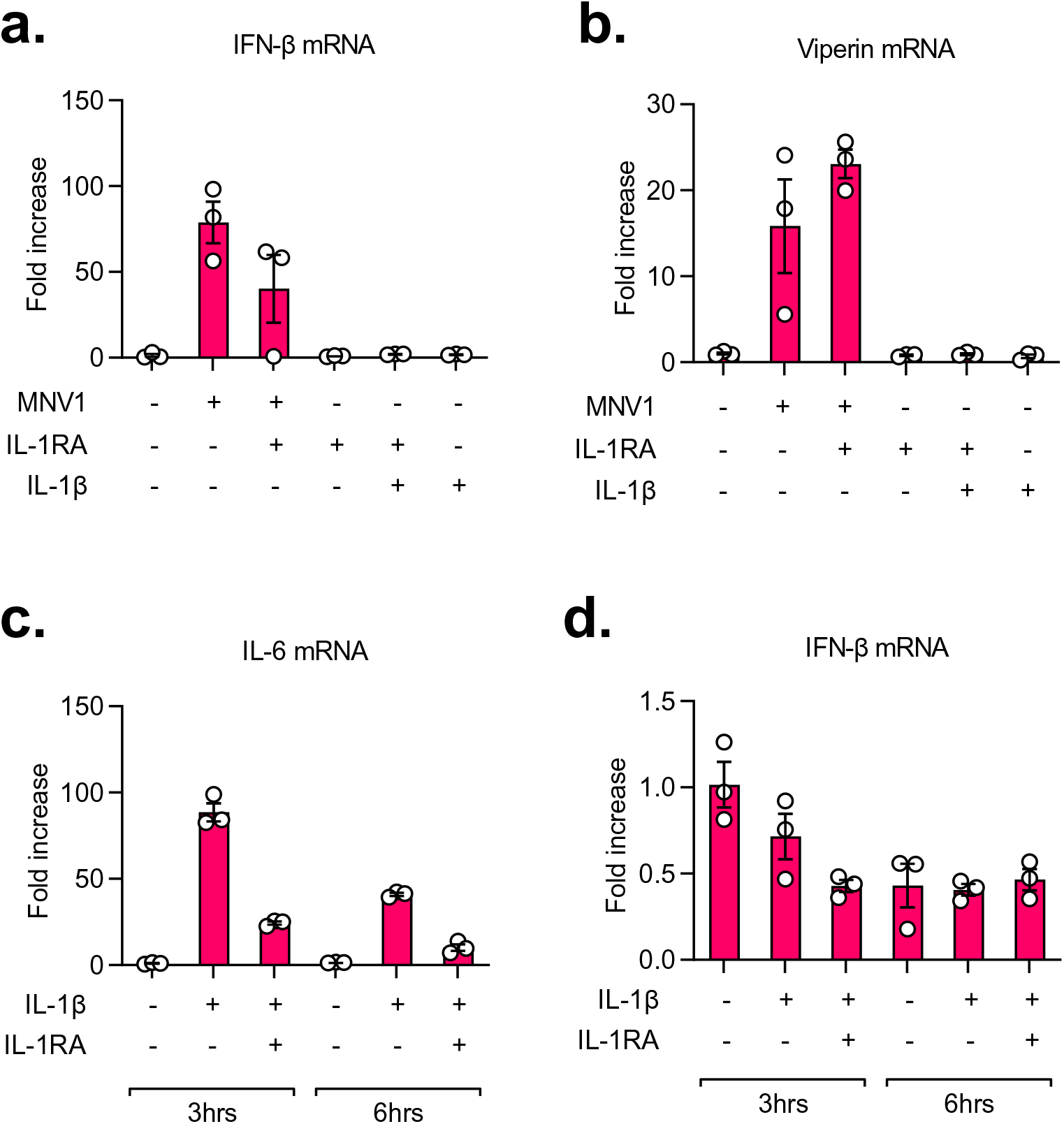
Inhibition of IL-1β signalling did not inhibit IFN responses in MNV-infected RAW264.7 cells, and IL-1β treatment failed to induce IFN-β in A549 cells. (a) and (b) RAW264.7 cells were either mock-treated or treated with 100ng/ml of recombinant IL-1RA for 30 minutes before infection with MNV1 at an MOI of 5, or treatment with 10ng of recombinant mouse IL-1β. The cells were harvested at 12h post infected and analysed by RT-qPCR. Data is presented relative to control and normalised to Gapdh (c) and (d) A549 cells were either mock-treated or treated with 100ng/ml of recombinant IL-1RA for 30 minutes before treatment with 10ng of recombinant human IL-1β. The cells were harvested at the indicated timepoints and analysed by RT-qPCR. Data is presented relative to control and normalised to β-actin.

**Figure S3.**
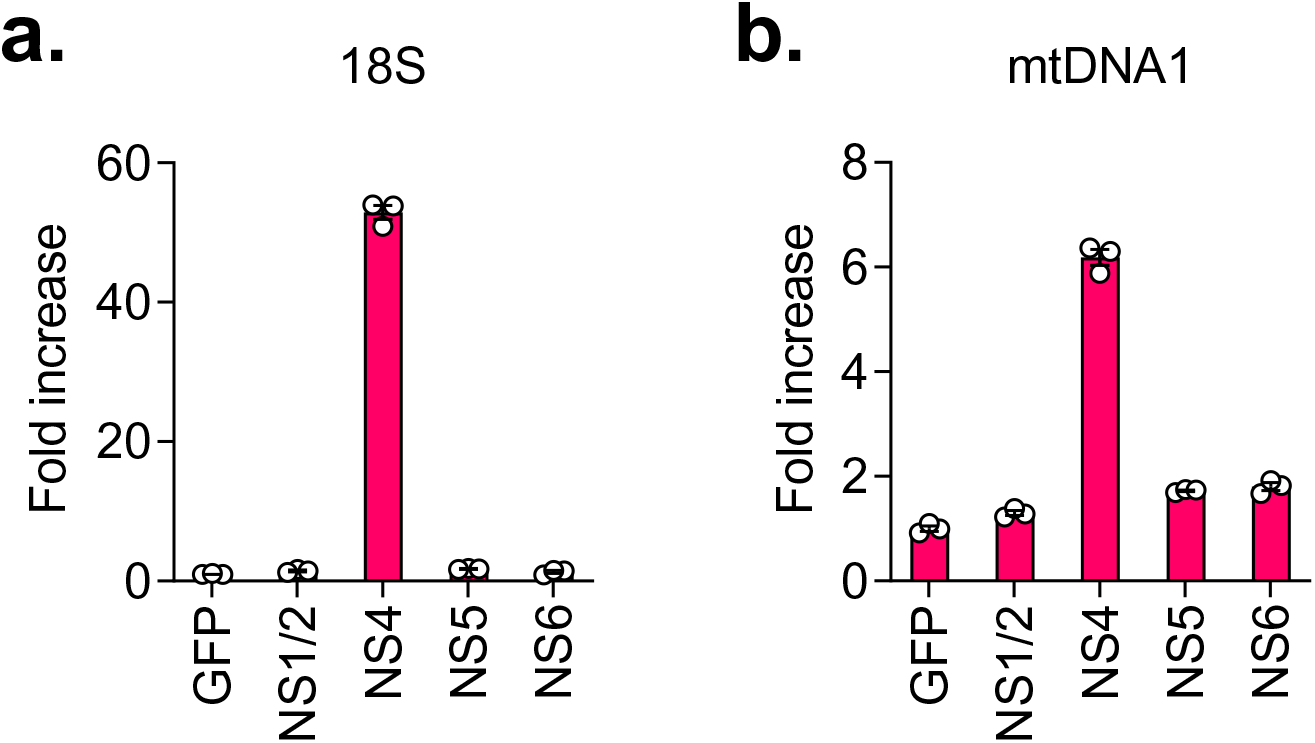
NS4, but not NS1-2 or NS6 (VPg), is sufficient for leakage of genomic and mitochondrial DNA in cell lines. (a) and (b) HEK293T-CD300lf cells were transfected with Flag-tagged construct plasmids of the indicated proteins. The cells were harvested 24h post transfection and assessed for cytosolic DNA by qPCR. Data is presented relative to GFP and normalised to their corresponding whole cell fractions.

